# Role of the NuRD complex and altered proteostasis in cancer cell quiescence

**DOI:** 10.1101/2025.02.10.637435

**Authors:** Qi Jiang, Michelle Ertel, Austin Arrigo, Sara Sannino, Jennifer L. Goeckeler-Fried, April Sagan, Betsy Ann Varghese, Daniel D. Brown, Wayne Stallaert, Adrian Lee, Amanda M. Clark, Jeffrey L. Brodsky, Hatice U. Osmanbeyoglu, Ronald J. Buckanovich

## Abstract

Cytotoxic chemotherapy remains the primary treatment for ovarian cancer (OvCa). Development of chemoresistance typically results in patient death within two years. As such, understanding chemoresistance is critical. One underexplored mechanism of chemotherapy resistance is quiescence. Quiescent cells, which have reversibly exited the cell cycle, are refractory to most chemotherapies which primarily target rapidly proliferating cells. Here, we report that CHD4 and MBD3, components of the nucleosome remodeling and deacetylase (NuRD) complex, are downregulated in quiescent OvCa cells (qOvCa). Indicating a direct role for NuRD complex downregulation in the induction of quiescence, either CHD4 or MBD3 knockdown or histone deacetylase inhibitors (HDACi), such as vorinostat, induce quiescence in OvCa cells. RNA-Seq analysis of HDACi-treated cells confirmed expression changes consistent with induction of quiescence. We also find that both primary qOvCa and vorinostat-induced qOvCa demonstrate altered proteostasis, including increased proteasome activity and autophagy, and combination therapy of HDACi and proteasome inhibitors or autophagy inhibitors demonstrated profound synergistic death of OvCa cells. Finally, we overlapped RNA-Seq signatures from quiescent ovarian cancer cells with genes essential for quiescence in yeast to identify a “quiescent cell core signature.” This core quiescent cell signature appeared to be conserved across multiple cancer types, suggesting new therapeutic targets.

## Introduction

Worldwide, ovarian cancer (OvCa) remains the most lethal gynecological cancer, with an extremely high mortality-to-incidence ratio [1]. An estimated 324,398 new cases of OvCa were diagnosed worldwide in 2022 [1]. Though most of these patients will have had an initial response to surgery and chemotherapy, 70% of patients will relapse and develop chemotherapy-resistant disease [2]. Despite the progress made on targeted and immune therapies, 206,839 patients succumbed to OvCa worldwide in 2022 [1]. Understanding the mechanism of chemotherapy resistance and seeking out potential methods to overcome resistance is essential for improving OvCa patient prognosis.

An important mechanism contributing to the development of chemotherapy resistance is cellular quiescence. Quiescent cells reversibly exit the cell cycle and stop proliferating and are thus refractory to standard chemotherapies, which preferentially target rapidly proliferating cells [3, 4]. With the appropriate signals, quiescent cells can re-enter the cell cycle and drive cancer recurrence. Understanding the factors that regulate quiescence could lead to better treatment approaches to prevent chemotherapy resistance.

Our greatest understanding of quiescent cell biology comes from studies of yeast as well as adult stem cells. Quiescent cells demonstrate epigenetic changes and changes in chromatin remodeling [5–7]. Epigenetic changes are associated with reduced ribosome biosynthesis, altered RNA processing, and altered protein translation [8, 9]. Furthermore, in the yeast *Saccharomyces cerevisiae*, the ubiquitin-proteasome system (UPS)—another key regulator of protein homeostasis (proteostasis)—also appears to be a critical regulator of the balance between proliferation and quiescence; for example, the proteasome assembly factor, Spg5, is induced in quiescent cells and is essential for yeast survival [10]. Notably, proteostasis is also critical for cancer cell survival [11].

While less studied, what is known of quiescence in cancer cells mirrors that which has been observed in yeast and mammalian adult stem cells. In various cancer types, quiescent cancer cells have been described as a subset of cancer stem-like cells (CSCs) and are linked to chemoresistance [12–18]. Yet, the contribution of quiescence in OvCa is ill-defined. However, studies in patient-derived xenograft (PDX) models demonstrated that OvCa cells that survived chemotherapy were primarily Ki-67 negative and enriched for expression of the CSC markers ALDH and CD133 [19]. In addition, Gao et. al. identified a similar population of slow-cycling, CD24+, chemotherapy-resistant OvCa cells that drive cancer recurrence in primary human tumors [20]. We recently reported that the NFATC4 transcription factor is activated by chemotherapy and can drive a chemotherapy-resistant quiescent cell state [21]. Furthermore, we found that, in response to chemotherapy, quiescent cells secrete follistatin to induce growth inhibition and chemotherapy resistance in neighboring cells. As such, follistatin knockout increased chemosensitivity and reduced chemotherapy resistance *in vitro* and *in vivo*.

To better define qOvCa cells, we recently performed single-cell RNA sequencing analysis on isolated primary qOvCa cells [22]. This analysis indicated that MBD3 and CHD4, two major components of the epigenetic regulatory nucleosome remodeling and deacetylase (NuRD) complex, are downregulated in qOvCa. While the NuRD complex has been linked to tumorigenesis, its role in quiescence remains relatively unexplored. In this study, we find that inhibition of NuRD complex activity, either by knockdown of MBD3/CHD4 or inhibition of the histone deacetylase activity of the NuRD complex with histone deacetylase inhibitors (HDACi), alters the expression of numerous cell cycle regulators and drives OvCa cells into a dense quiescent state. Furthermore, both primary quiescence and HDACi-induced quiescence were associated with altered proteostasis, including upregulation of proteasome activity and autophagy-associated proteins. Suggesting an important role for these alterations in proteostasis, we found that HDACi-induced quiescent cells are sensitized to drugs, such as proteasome inhibitors and autophagy inhibitors, which alter proteostasis. Finally, we overlaid multiple qOvCa RNA-Seq data sets with yeast essential quiescence genes to further define a quiescent cancer cell signature. This effort yielded a core quiescence signature which is reflected across multiple tumor types.

## Results

### NuRD complex components MBD3 and CHD4 are downregulated in quiescent OvCa

We recently performed single-cell RNA sequencing of primary qOvCa [22]. This analysis indicated qOvCa cells had downregulated mRNAs corresponding to two components of the NuRD complex, MBD3 and CHD4 [6]. Western blot analysis in control OvCa cells and serum starvation-driven qOvCa cells confirmed downregulation of both MBD3 and CHD4 protein in three OvCa cell lines (Fig. 1A). To determine whether downregulation of MBD3 and CHD4 was a result, or potential driver, of quiescence, we next performed siRNA and shRNA knockdown studies. The efficiency of knockdown was validated by RT-qPCR and Western blot (Fig. S1A-S1D). The knockdown of either MBD3 or CHD4 resulted in a decrease in cell proliferation (Fig. 1B, Fig. S1E) and no change in cell viability as measured by an apoptosis assay (Fig. 1C, Fig S1F). Furthermore, cell cycle analysis using the Fucci fluorescent cell cycle reporter system [23] demonstrated that the knockdown of MBD3 and CHD4 was associated with an increase in p27Kip/CDT1-positive cells/cells in the G0 phase of the cell cycle (Fig. 1D). These findings implied that decreased expression of MBD3 and/or CHD4 can induce cellular quiescence in OvCa.

**Figure 1.**
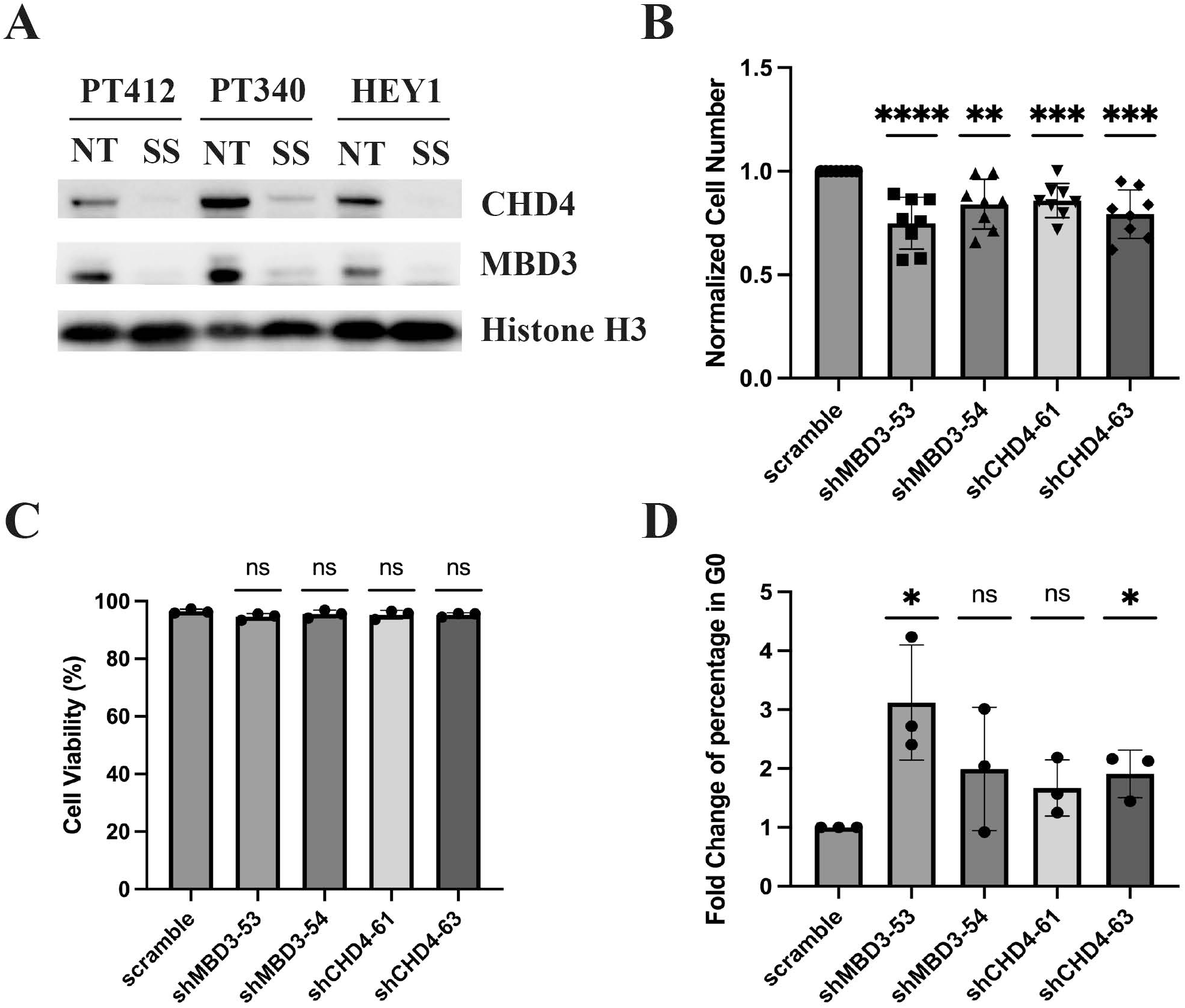
Knockdown of MBD3/CHD4 drives quiescence in OvCa cells. **A.** Western blot analysis of MBD3 and CHD4 in control and serum starvation-induced quiescence in PT412, PT340, and Hey1 cell lines. **B.** Cell numbers in PT412 cells with scrambled control, MBD3, or CHD4 shRNAs. **C.** Cell viability of PT412 cells transfected with scrambled, MBD3, and CHD4 shRNA and assessed via PI/Annexin-V apoptosis assay. **D.** Summary of cells determined to be in G0 using flow cytometry of Fucci reporter (CDT1(30/120)-mcherry/p27K^-^-mVenus)-labelled Hey1 cell line treated with scrambled control, MBD3, and CHD4 shRNA.

### Inhibition of the NuRD complex by HDAC inhibitors drives a dense quiescent state in OvCa cells

Because the redundancy of MBD/CHD family proteins may reduce the impact of knockdown, we next evaluated whether pharmacologic inhibition of the NuRD complex using HDAC inhibitors (HDACi) would induce quiescence. We treated OvCa cells with the pan-HDAC inhibitors vorinostat (SAHA), belinostat, or valproic acid (VPA), or the HDAC1, 2, 3, 11 selective inhibitor mocetinostat for 48 hours.

Consistent with the induction of quiescence, HDACi-treated cells showed a significant reduction in cell number with minimal impact on cell death (Fig. 2A-B). Cell cycle analysis, using the Fucci cell cycle reporters (p27-mVenus and CDT1-mcherry), suggested each of the HDACi induce a dense quiescent state, with >90% of cells arrested in the G0 phase of the cell cycle (Fig. 2C). To confirm this finding, vital dye (cell trace violet-CTV) was used to label OvCa cells used to track cellular proliferation, and the cells were then treated with vorinostat. Flow cytometry demonstrated that vorinostat treatment significantly increases the percentage of CTV-bright (i.e., non-dividing) cells compared to DMSO control cells (Fig. 2D). To determine whether vorinostat-induced cell cycle arrest is reversible, we performed real-time imaging of OvCa cells. Replicated groups of cells were treated with (i) DMSO, (ii) DMSO for 24 hours (allowing cells to reach linear growth) followed by vorinostat, or (iii) DMSO for 24 hours, then vorinostat for 48hrs, and then vorinostat washed out and DMSO was re-applied. While maintenance of vorinostat- restricted cellular proliferation compared to DMSO control, indicating reversibility of cellular arrest, vorinostat washout allowed cells to resume proliferation within ∼24 hours in multiple OvCa cell lines (Figure 2E, Figure S2).

**Figure 2.**
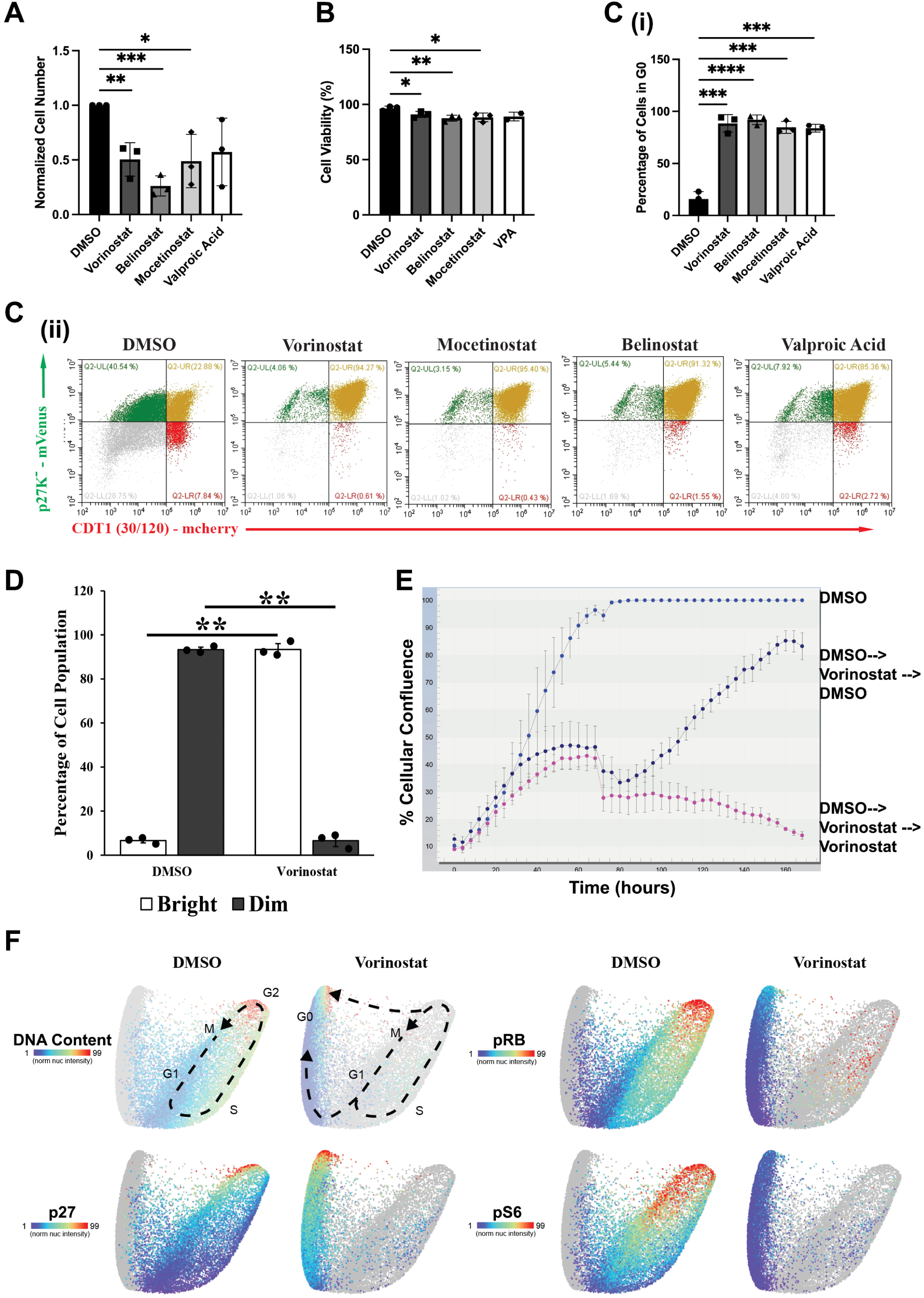
Pharmacologic inhibition of NuRD complex induces quiescence in OvCa cells. **A.** Normalized cell number in DMSO-, vorinostat-, mocetinostat-, belinostat-, and VPA-treated OvCa cell lines (PT412, PT340, Hey1). **B**. Cell viability in DMSO-, vorinostat-, mocetinostat-, and belinostat-treated PT412, PT340, and Hey1 and VPA-treated PT412 and Hey1. **C** Summary (i) and (ii) representative flow cytometry plots of Fucci reporter-labeled Hey1 cell line treated with DMSO, vorinostat, mocetinostat, and belinostat. **D**. Summary of flow cytometry assessment of cell trace violet (CTV)-labeled retention/dilution of OvCa cells treated with or without vorinostat for 5 days. Percentage of Dim (proliferating) or Bright (quiescent) cells in total cell population. **E**. Real-time cell proliferation analysis of Hey1 cells treated with (1) DMSO, (2) vorinostat 72h followed by DMSO (vorinostat**→**DMSO), (3) vorinostat. **F**. Cell cycle map of DMSO- and vorinostat-treated PT340 cells. (Dotted lines indicate proliferative and arrest trajectories.). Each experiment was independently repeated three times, with p values calculated using ANOVA, comparing means between independent studies.

Next, to further confirm induction of quiescence, we used multispectral iterative immunofluorescence to evaluate the expression for 28 cell cycle effectors, including CDKs, cyclins, p21, p27, phospho-Rb, and DNA content in DMSO control and vorinostat-treated OvCa cells. The intensity of phospho-Rb, p27, and other cell cycle regulators in DMSO treated cells was then used to define a cell cycle map, as previously described [24–28]. Vorinostat-treated cells demonstrated a staining pattern with the majority of cells in G0 and pre-G0 state, which included a reduction in nuclear p27, pRB, and pS6 (Fig. 2F, Fig. S3).

Together, these data indicate that loss of NuRD complex function, via either genetic downregulation of CHD4 or MBD3 or pharmacologic inhibition by HDAC inhibitors, induces OvCa cellular quiescence.

### Vorinostat-induced quiescent cells have transcriptional changes similar to primary quiescent cells

To examine the transcription changes associated with vorinostat-induced quiescence in OvCa cells, we performed bulk RNA sequencing. Two OvCa cell lines were treated with DMSO or vorinostat for 48 hours, and the total RNA was extracted and used for bulk RNA sequencing (n=3/treatment group). The principal component analysis (PCA) plot showed that the triplicate samples were consistent and that the treatment group had a consistent treatment effect (Fig. S4). Consistent with this, a heatmap of the top 15,000 expressed genes also showed samples clustered by treatment instead of cell line (Fig. 3A). An analysis (log fold change (LFC) >1 or < -1, p-adjust <0.05) of differentially expressed genes (DEG) revealed 5543 DEGs in PT340 and 6105 DEGs in PT412 cells, with 4137 DEGs overlapping in the two cell lines (Fig. 3B) [29].

**Figure 3.**
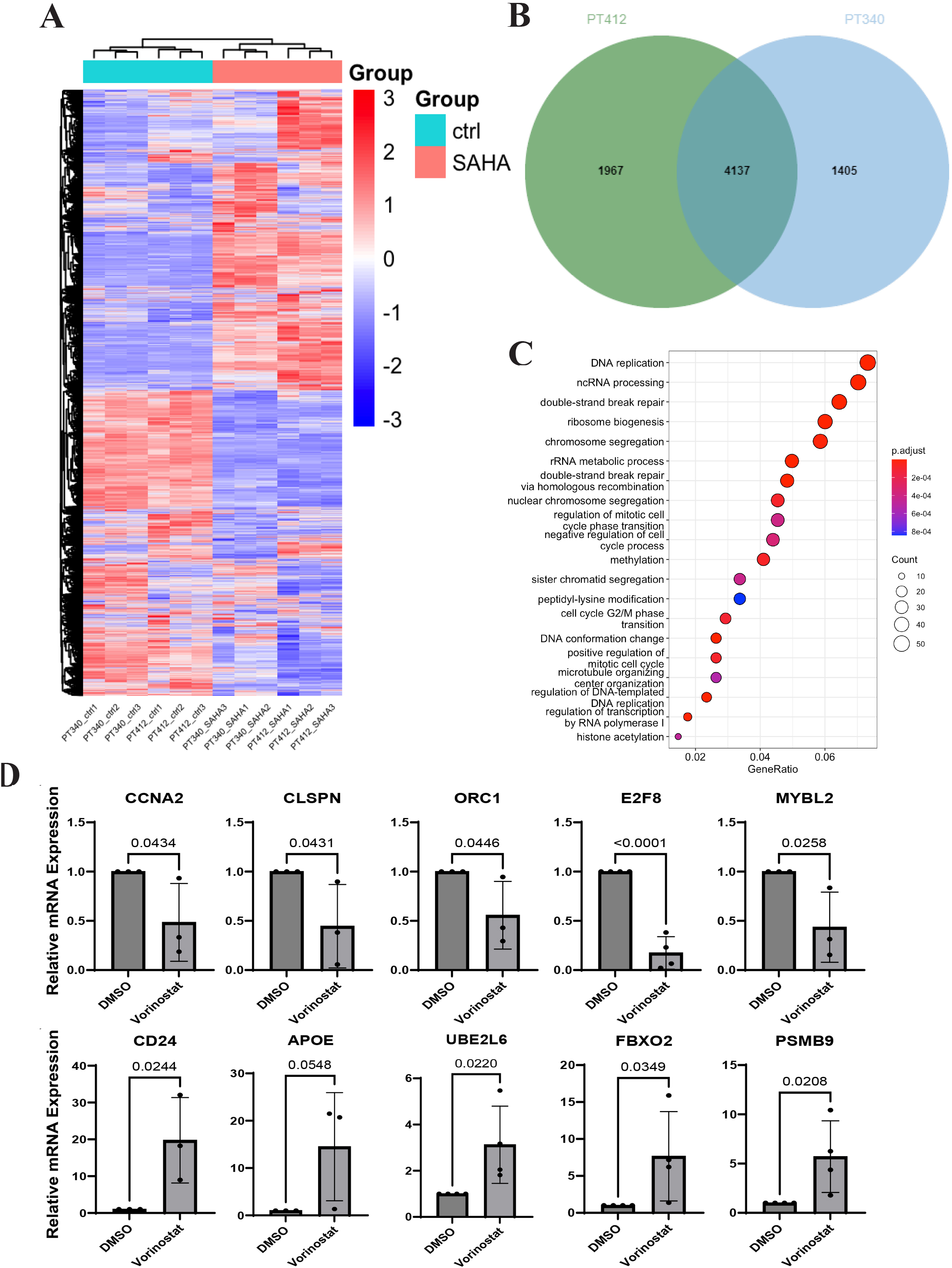
RNA sequencing of vorinostat-treated cells shows characteristics of quiescent cells. **A.** Heatmap of RNA-seq of top 15,000 genes in control and vorinostat-treated cells in PT340 and PT412. **B.** Venn diagram showing numbers of overlapped DEGs in vorinostat-treated PT340 and PT412. **C.** GO biological process analysis for genes downregulated (LFC<1.0 and p.adj<0.05) in vorinostat-treated PT340 and PT412 cells, showing top 20 pathways. **D.** RT-qPCR evaluation of expression of the indicated genes in control and vorinostat-treated OvCa cell lines (PT340, PT412, A2780).

Gene oncology (GO) biological process analysis of overlapped downregulated DEGs demonstrated alterations in ribosome biogenesis, RNA processing, DNA replication, cell cycle phase transition, cell division, and DNA conformation, all of which are consistent with the characteristics of quiescent cells (Fig. 3C) [30]. In addition to expression changes in numerous cell cycle-related genes, ubiquitin-proteasomal system (UPS)-related genes were also differentially expressed in vorinostat-treated cells, as compared to controls, across both cell lines. RT-qPCR analysis of cell cycle factors (CCNA2, CLSPN, E2F8, MYBL2, ORC1), cancer progression marker APOE [31], cancer stem cell marker CD24 [32, 33], and the UPS factors (UBE2L6, FBXO2, PSMB9) confirmed the RNA-Seq data (Fig. 3D). These results are also consistent with the iterative immunofluorescent cell cycle mapping, which showed CCNA2, CCNB2, and CCND1 to be downregulated at the protein level (Fig. S3). Combined, these data indicate that vorinostat treatment induces quiescence in OvCa cells.

### Targeting proteostasis induces quiescent OvCa cell death

Many of the altered pathways identified in our RNA-Seq studies, including ribosome biogenesis, RNA processing, translation initiation and the UPS, play critical roles in regulating proteostasis [34]. To study whether proteostasis is essential for quiescent OvCa cells survival, we screened five therapeutics that disrupt proteostasis: carfilzomib (a proteasome inhibitor), tanespimycin (an HSP90 inhibitor), hydroxychloroquine (an autophagy inhibitor), pidnarulex (an rRNA synthesis inhibitor), and seclidemstat (an LSD-1 inhibitor) (Table 1). We tested each drug, alone and in combination, with vorinostat.

**Table 1.**
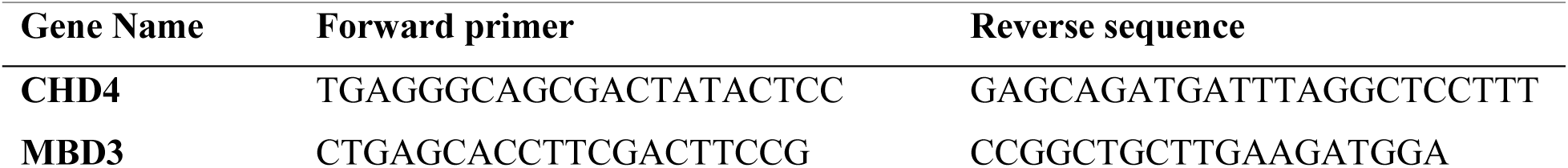
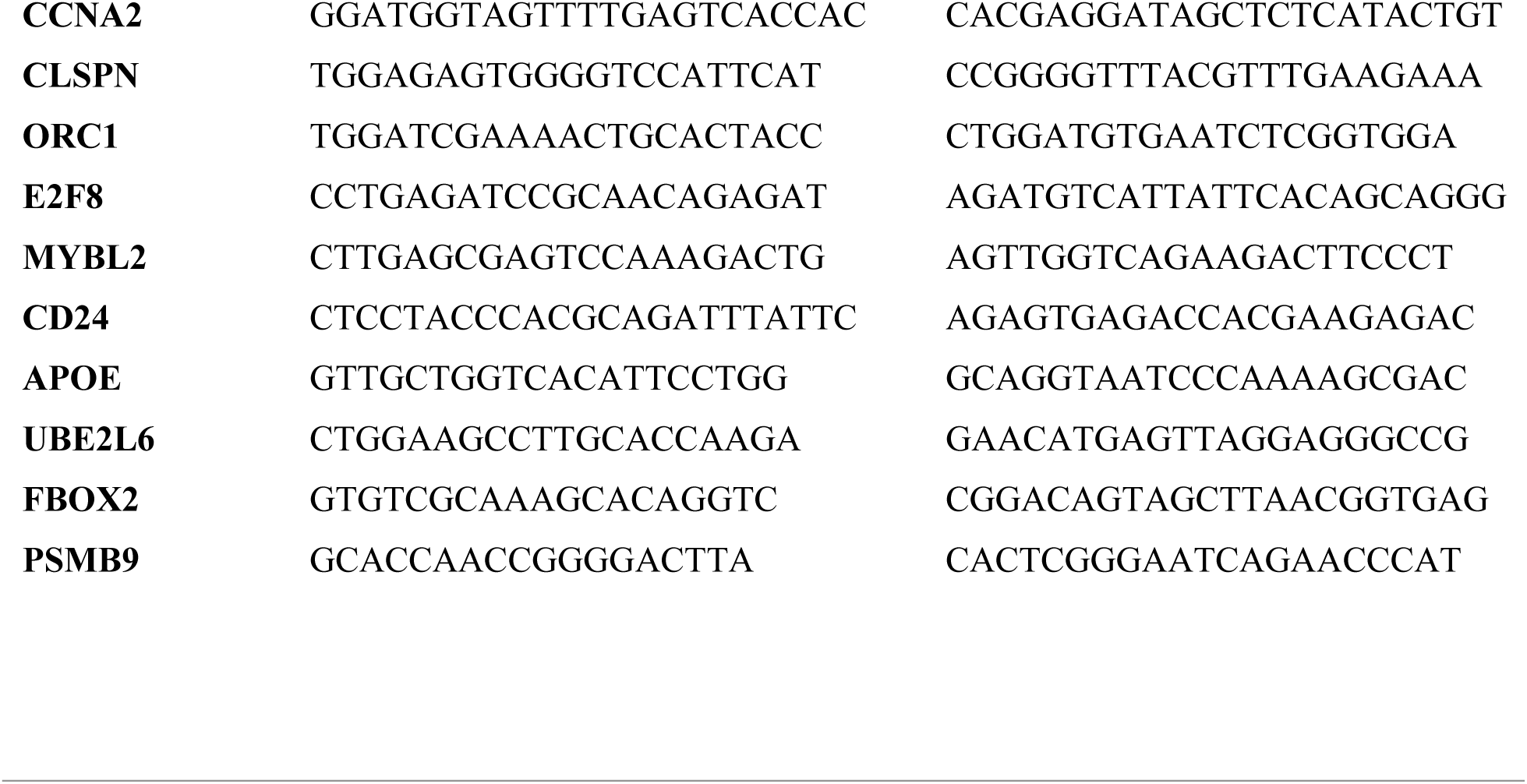
Primers sequences.

Treatment with tanespimycin, carfilzomib, hydroxychloroquine, and seclidemstat as single agents demonstrated a 20-30% reduction in cell numbers, while single-agent pidnarulex demonstrated a ∼75% reduction in cell numbers (Fig. 4A, Fig. S5A). Each drug, as a single agent, had little or no impact on cell viability. In contrast, when combined with vorinostat to induce quiescence, carfilzomib, hydroxychloroquine, and pidnarulex resulted a significant reduction in cell viability (Fig. 4B, fig. S5B).

**Figure 4.**
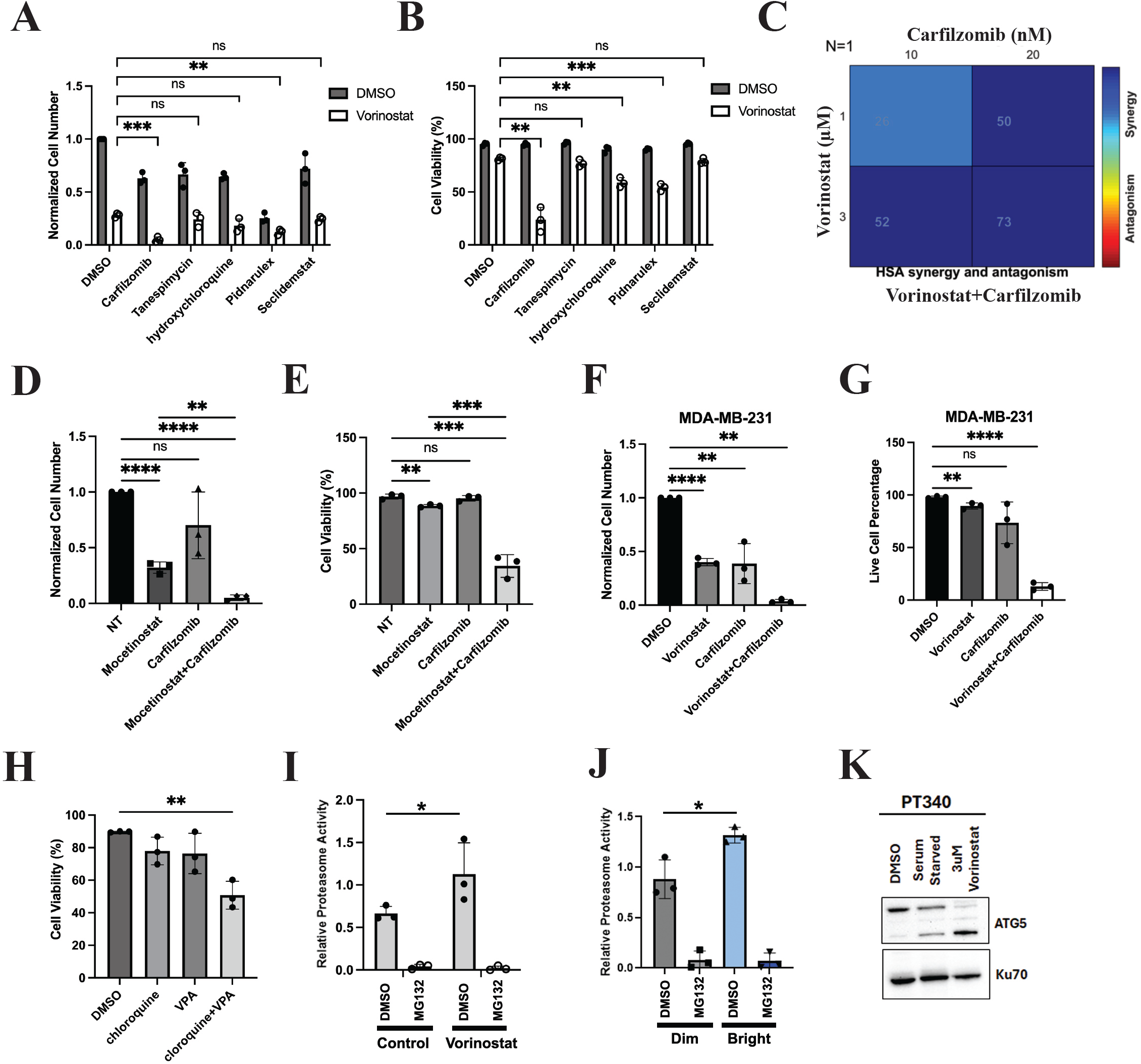
qOvCa cells have altered proteostasis, and targeting proteasome induces quiescent cancer cell death. **A** and **B.** Normalized cell number and viability of PT412 cells treated with DMSO or vorinostat (SAHA) and the indicated compounds. **C.** HSA synergy heat map for vorinostat and carfilzomib. **D.** Normalized cell number of DMSO, mocetinostat, carfilzomib single-drug treatment, and combination treatment in PT340. **E.** Cell viability of DMSO, mocetinostat, carfilzomib single-drug treatment, and combination treatment in PT412. **F.** Normalized cell number of DMSO, vorinostat, carfilzomib single-drug treatment, and combination treatment in MDA-MB-231. **G.** Cell viability of DMSO, vorinostat, carfilzomib single-drug treatment, and combination treatment in MDA-MB-231. **H.** Cell viability of DMSO, chloroquine, and VPA single-drug treatment, and combination treatment across OvCa cell lines (PT340, PT412, A2780). **I.** Proteasome activity in control and vorinostat-treated OvCa cell lines (PT340, PT412, and A2780). To confirm specificity, proteasome activity was suppressed by the proteasome inhibitor MG132. **J.** Proteasome activity in primary quiescent, and proliferating (vital dye bright and dim, respectively) PT340 cells. **K.** Western blotting probed ATG5 in DMSO-, serum starvation-, and vorinostat-treated PT340 cells. Cell number/viability and proteosome studies were completed in triplicate, with p values calculated using ANOVA, comparing means between independent studies.

Among the combination therapies, the vorinostat and carfilzomib combination resulted in the most significant cell death. This was striking as neither drug alone induced significant cell death. Indeed, synergy calculation, using Combenefit [35] with the HSA model, demonstrated that vorinostat and carfilzomib had profound synergistic effects (synergy index 25-73) on OvCa cell viability (Fig. 4C). Confirming the importance of HDAC inhibition on the effect of combination therapy, mocetinostat (Fig. 4D-E, Fig. S5C) and belinostat (Fig. S5D-E) similarly showed synergistic effects with carfilzomib in OvCa cell lines. To determine if this was ovarian cancer specific, we repeated these studies in the breast cancer cell line MDA-MB-231. Similar to the studies in OvCa lines, HDAC inhibitor treatment also decreased MDA-MB-231 cell growth while having minimal impact on cell viability, and combination therapy with vorinostat and carfilzomib significantly increased cell death (Figure 4F-G).

To further confirm a critical role for proteostasis in HDACi-induced quiescent cells, we performed similar combination studies of HDACi with the autophagy inhibitor chloroquine. For these studies we chose to use the HDACi valproic acid as it is significantly cheaper to use for patients and, thus, combined with chloroquine, could provide an easier path to the clinic. While neither agent demonstrated significant single-agent toxicity, combination of chloroquine and VPA resulted in a 50% reduction in cell viability across three OvCa cell lines (PT340, PT412, and A2780; Figure 4H).

To determine why vorinostat-induced quiescent cells are sensitive to proteasome inhibitors, we measured the proteasome activity in control and vorinostat-treated cells. We therefore monitored the liberation of 7- amino-4-methylcoumarin from the synthetic proteasome substrate Suc-LLVY-AMC [36]. The data shown in Figure 4I indicate that vorinostat-treated OvCa cells had a trend of increased proteasome activity compared to the DMSO control. To show that this relationship mirrored proteasome activity in primary quiescent cells, we similarly evaluated proteasome activity in proliferating and quiescent primary cells based on CTV vital dye stain. CTV-bright cells/non-proliferating demonstrated an increase in proteasome activity proportional to the increase in proteasome activity in vorinostat-treated cells (Figure 4J).

Similarly, analysis of the autophagy marker ATG5 revealed the upregulation of autophagy in both serum starvation and vorinostat-induced quiescent cells (Figure 4K). Combined, our data suggest that the proteasome and autophagy are upregulated in OvCa quiescent cells, which may be essential to support cell viability.

### HDACi and a proteostasis inhibitor act synergistically in therapeutic tumor models

We next evaluated the therapeutic effect of single-agent and combination therapy *ex vivo* and *in vivo*. First, two patient-derived triple-negative breast cancer organoid models (IPM-BO-085 and IPM-BO-139) were treated with DMSO, vorinostat, or carfilzomib, or a combination of the two drugs. The viable cell number was then assessed with the CellTiter-Glo 3D cell viability assay. Single-agent vorinostat had no effect in IPM-BO-085 organoids but reduced growth by approximately 50% in IPM-BO-139 organoids. In contrast, single-agent carfilzomib reduced growth by 20% and 60%, respectively, in the two models.

However, a combination of vorinostat and carfilzomib treatment significantly reduced organoid viability in both models, with an 85-90% reduction in viable cell number (Figure 5A).

**Figure 5.**
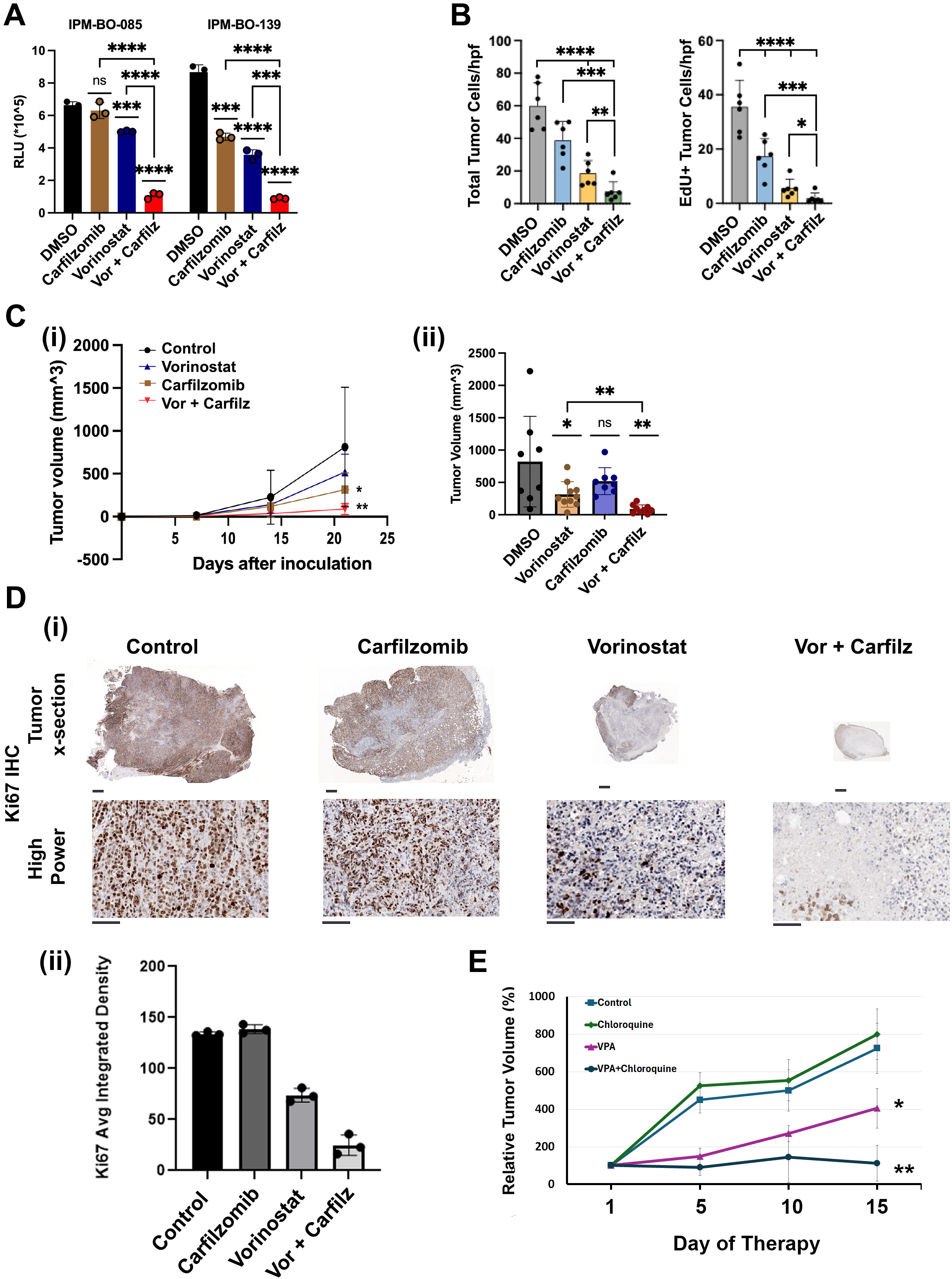
Combined targeting of proteasome and NuRD complex delays tumor growth *in vivo*. **A**. Cell viability, as assessed by CellTiter-Glo 3D, of patient-derived triple-negative breast cancer organoids treated with DMSO, vorinostat, carfilzomib single-drug treatment, and combination treatment. **B** Total and EdU+ viable MDA-MB-231 breast cancer cells co-cultured with hepatocytes and treated with the indicated therapies. Data are pooled results from replicate experiments compared with students t-test. **C.** Tumor growth curves and final tumor volumes from NSG mice injected with PT340 cells subcutaneously. Mice were treated with vehicle control, vorinostat, carfilzomib single-drug treatment, and combination drugs 5 days after inoculation. **D**. (i) Low-magnification (whole tumor cross-section) and high-power images of Ki67 staining indicate treatment groups, and (ii) quantification of Ki67 average integrated density from 10 high-power sections of three tumors per treatment group. **E**. Tumor volume curve of NSG mice injected with PT340 cells subcutaneously. Mice were treated with vehicle control, valproic acid (VPA) or chloroquine single-drug treatment, and combination VPA and chloroquine 5 days after inoculation.

Next, we evaluated the therapeutic response in an *ex vivo* 2D metastasis model, using a co-culture of primary murine hepatocytes and RFP-labeled MDA-MB-231 cells. Cells were co-cultured for 24 hours and then treated with DMSO, vorinostat (1.5uM), carfilzomib (10nM), or the combination for 72 hours before total number of viable cancer cells and EdU+ cancer cells were assessed. Single-agent vorinostat treatment was associated with a significant reduction in total and EdU+ cancer cell numbers. Mirroring the results shown above, combination therapy was again associated with significant reduction in total cell numbers, and near complete loss of EdU+ cells (Fig 5B).

We also evaluated single and combination drug therapy *in vivo*, using a subcutaneous ovarian PDX model. PT340 cells were injected into the flanks of NSG mice, and daily drug treatment were started 5 days after inoculation. Single-agent carfilzomib did not significantly impact tumor growth. Consistent with an induction of quiescence, single-agent vorinostat significantly slowed tumor growth, while— consistent with tumor cell killing—combination therapy with vorinostat and carfilzomib was highly effective at limiting tumor growth, with tumors undetectable in two of nine mice (Fig 5C).

Immunohistological analysis of tumors for Ki67 expression demonstrated that >90% of cells in the control and carfilzomib treatment groups were proliferative, with less than 50% proliferative in the reviewable vorinostat-treated tumors, while dual therapy-treated tumors showed a profound reduction in Ki67 stand-in and large areas of necrosis (Fig 5D).

To further confirm the therapeutic benefits of combination quiescence-inducing/proteostasis-targeting drugs, we repeated the *in vivo* study with valproic acid and chloroquine. Similar to the study above, single-agent chloroquine (proteostasis-targeting agent) had no treatment effect, while single-agent valproic acid (quiescence-inducing agent) significantly delayed tumor growth. However, combination therapy again prevented tumor growth and induced mild tumor regression (Fig. 5E).

### Identification of a core quiescence signature

Our studies, and those of others, indicate that quiescent cells have unique attributes, including altered proteostasis, which can be exploited for therapeutic purposes. To better define the expression changes and possible therapeutic targets in quiescent cells, we used existing data sets to identify a universal cancer “quiescent cell core signature.” To this end, we overlaid our existing RNA-Seq datasets with two different quiescent cell RNA signatures identified in *Schizosaccharomyces*, a well-established model of quiescence; specifically, Klosinska et al. reported the transcriptional profiles of nutrient depletion-induced quiescence [37], and Sajiki et. al. reported genes essential for the mitotic competence for *Schizosaccharomyces* cells to exit quiescence [38]. We overlaid these datasets with human orthologs identified after RNA sequencing of (i) vorinostat-induced quiescent cells, and (ii) the recently reported MRTFA inhibitor-induced quiescent cell signature [22]. Based on our analysis, we identified 25 genes, including upregulated genes (e.g., SLC7A8, WIPI1, and SERINC2) and downregulated genes (e.g., CCNF, CCNB1, HSPA14, and IRAK1), which we suggest represents a conserved quiescent cell core signature (Figure 6A, 6B). To validate the core signature across cancer types, we performed RT-qPCR in DMSO-, serum starved-, vorinostat-, and CCG-257081-treated OvCa, lung cancer, and breast cancer cells. Consistent with the identification of a core signature, SERINC1, WIPI1, CCNF, CCNB1, and CCNA2 were similarly regulated across cancer types under different treatments (Figure 6C).

**Figure 6.**
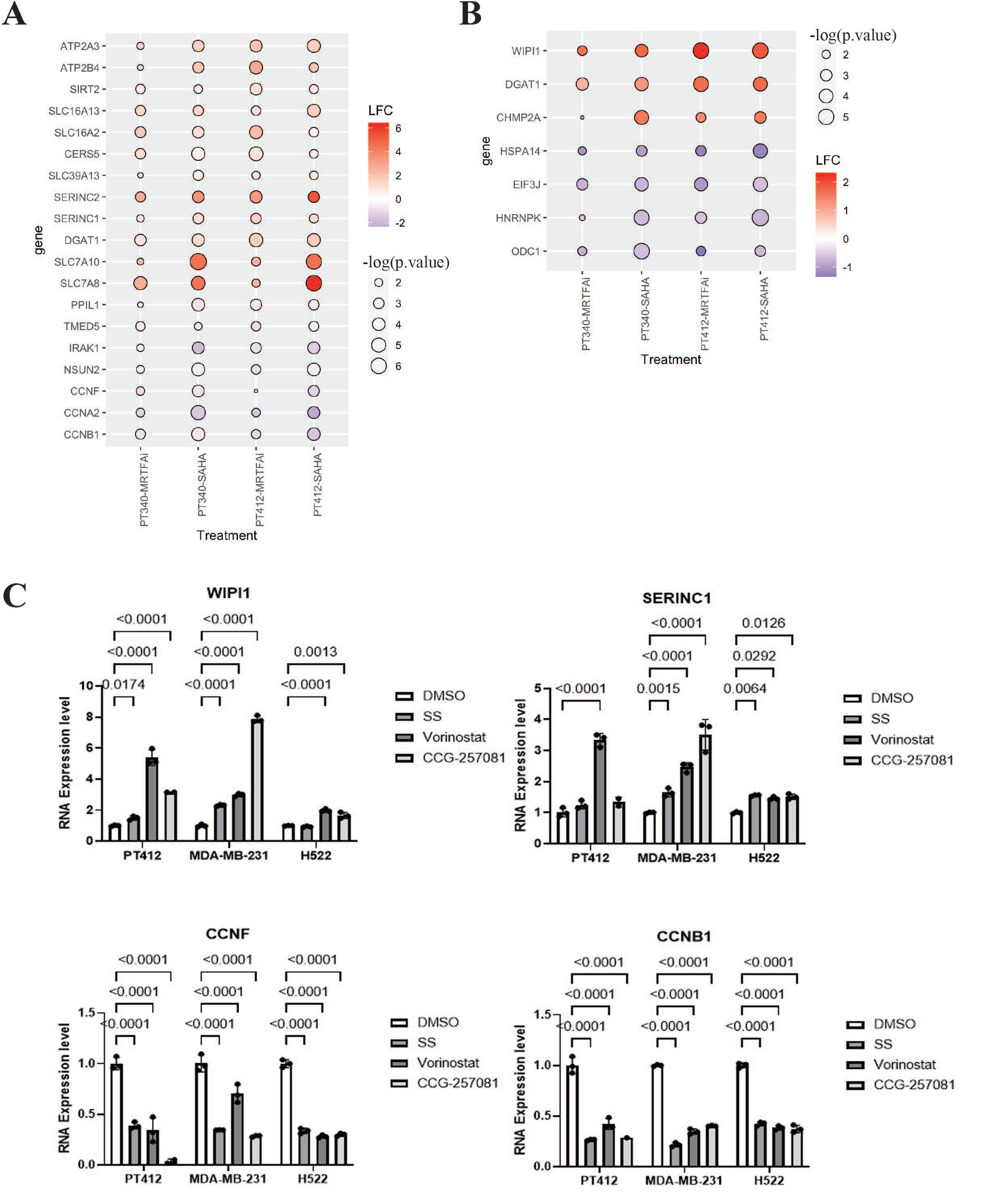
Core quiescence signature shared by *Schizosaccharomyces* and OvCa cells. **A.** Heat map of quiescence-essential genes in *Schizosaccharomyces* and differentially expressed genes from ovarian cancer cells treated with quiescence-inducing drugs (vorinostat and MRTFA inhibitor CCG081). Cutoffs in qOvCa cells are LFC>0.5 or <-0.5, p.value<0.05; Cutoff in *Schizosaccharomyces*: Quiescent effect>1, p.value<0.05. **B.** Heat map of *Schizosaccharomyces* mitotic competence-essential genes altered in differentially expressed genes from ovarian cancer cells treated with quiescence-inducing drugs. (Cutoffs in qOvCa cells are LFC>0.5 or <-0.5, p.value<0.05). **C.** Validation of quiescence core gene alternation in serum starvation-, vorinostat-, and CCG-257081-induced quiescence in PT412, MDA-MB-231, and H522 cell lines.

## Discussion

We report, here, a critical role for NuRD complex downregulation in induction of quiescence in OvCa. We also report that quiescent OvCa cells exhibit dependencies related to altered proteostasis, including increased proteasome activity. Importantly, the increase in proteasome activity appears to be an Achilles heel for quiescent cells. While bulk ovarian cancer cells are, in general, therapeutically resistant to proteasome inhibition, HDACi-induced quiescent cells are killed when a clinical proteasome inhibitor is added. Finally, by overlaying RNA-Seq data sets from quiescent cancer cells and yeast, we identify a conserve core quiescent cell signature. This signature was validated in quiescent cells across several cancer types.

### The NuRD complex and quiescence

We found that MBD3 and CHD4, two components of the NuRD complex, are downregulated in primary quiescent cells at both the RNA and protein level. Inhibition of the NuRD complex via either genetic knockdown or pharmacologic inhibition with HDACi can enforce a quiescent state in OvCa cells. However, the effect of knockdown was milder than that of pharmacologic inhibition. We speculate this is due to the redundancy of CHD/MBD family proteins. CHD3, CHD4, and CHD5 are three highly homologous proteins and can form distinct NuRD complexes that contain other components [39–41]. Since CHD5 is preferentially expressed in testis and the nervous system, CHD3 and CHD4 are more commonly present in the NuRD complex. [41, 42]. Studies revealed that CHD3-NuRD and CHD4-NuRD have different localization patterns and distinct regulated genes, but there are common target genes equally influenced by CHD3-NuRD and CHD4-NuRD [39]. Likewise, MBD2 and MBD3 have been reported to form mutually exclusive complexes with different functions but can also bind to the same promotor regions [43, 44]. Furthermore, evidence showed that, in the absence of CHD4, other components can form a stable complex with robust HDAC activity, indicating that CHD4 is a peripheral component in the NuRD complex [45]. Therefore, knockdown of a single CHD/MBD is insufficient to fully inhibit NuRD complex activity.

HDACi were found to strongly induce quiescence in OvCa cells, and indeed they are proven to have anti-tumor effects in multiple others cancers [46–48]. Vorinostat, which inhibits the activity of class I and II HDACs, was the first FDA-approved HDACi used for refractory cutaneous T-cell lymphoma [48, 49].

Consistent with our results, vorinostat treatment induced cell cycle arrest in breast cancer and in non-small cell lung cancer cell lines [50, 51]. Others have reported an effect in OvCa cells; however, the phase of cell cycle arrest is controversial [52–54].

Although vorinostat exhibits a satisfactory anti-proliferative effect in solid tumor cell lines, it has not demonstrated significant activity as a single agent in phase II clinical trials for refractory glioblastoma, breast cancer, colorectal cancer, non-small cell lung cancer, or epithelial ovarian cancer [55–57]. HDACi induction of quiescence would be consistent with these results; induction of quiescence would be expected to slow tumor growth or, at best, induce stable disease, but would be unable to reduce tumor volume. As most clinical trials have primary endpoints related to radiologic reduction in tumor volume, these trials were destined to fail. Our results suggest that use of HDACi as a maintenance therapy to delay recurrence would be a more efficacious approach. Furthermore, given that quiescent cells are typically chemotherapy resistant, our results further suggest that HDACi should not be combined with chemotherapy.

### Quiescence and proteostasis

Our studies are consistent with both primary quiescent cells and vorinostat-driven quiescent cells exhibiting enhance proteosome and autophagy activity. Our results are also consistent with other work indicating that vorinostat induces proteasome activity [58–62] and is in line with prior studies indicating that many proteosome-related pathways are altered in quiescent cells [7]. The upregulation of these proteostasis pathways indicates they may support quiescence. Indeed, the preclinical studies performed here indicate that proteasome activity is critical for quiescent cancer cell survival. As such, approaches targeting these pathways could help eradicate residual, chemotherapy-resistant, quiescent cancer cells and could have a profound impact on the outcome of cancer patients. The fact that similar treatment effects were also noted in breast cancer cells suggests that this is not a cancer type-specific affect. It is interesting to note that proteosome inhibitor therapy has proven most effective in multiple myeloma – a “smoldering,” more quiescent cancer type. Similarly, given that neurons are generally felt to be quiescent cells, it may not be coincidence that neurotoxicity is the major side effect of proteosome inhibitors.

Autophagy is another proteostasis-related pathway, and has been shown to be dysregulated in cancer stem cells [63]. We found that a key autophagy-associated protein is upregulated in qOvCa cells. Dual HDACi therapy and autophagy inhibition with hydroxychloroquine or chloroquine induced OvCa cell death *in vitro* and delayed tumor growth *in vivo*. Parallel to our finding, Rehman et al. revealed that a reversible drug-tolerant persister state, induced by CPT-11, is maintained by an upregulated autophagy pathway in colorectal cancer cells [64]. They demonstrated that a combination of the autophagy inhibitor chloroquine and CPT-11 robustly increased apoptosis.

### Quiescent cell core signature

These results and others indicate that quiescent cells may have a unique biology that can be exploited for therapeutic benefit. To identify additional targets in quiescent cancer cells, we identified genes in common between quiescent cancer cells and quiescent yeast. This core quiescent signature suggests several new therapeutic targets, including several solute carriers and cell cycle-related genes. In the yeast studies, some genes were identified in response to quiescence-driving stimuli, while others were necessary for “mitotic competency” or the ability to replicate after quiescence. We speculate that these mitotic competency genes may include genes that help drive quiescent cells back into the cell cycle. Further studies will be needed to confirm this hypothesis.

In summary, this study provides insight on the NuRD complex’s function in regulating cancer cell quiescence. By inhibiting NuRD complex activity, OvCa cells showed a decreased proliferation rate but with high cell viability and ability to regrow rapidly after drug removal. Increased proteasome activity helps qOvCa cells maintain proteostasis and survive under stress. With combinatorial inhibition of NuRD complex and proteasome activity, most OvCa cells were killed *in vitro*. The combination therapy of vorinostat and carfilzomib *in vivo* showed profound effects in delaying tumor growth, which indicates a possible new therapeutic method for OvCa patients.

### Materials and Methods Cell lines and culture

PT340 cells were derived from an abdominal metastasis of a mixed high-grade serous/clear cell ovarian cancer [65]. PT412 was a patient-derived, stage III, abdominal metastasis, high-grade serous ovarian cancer cell line from Dr. Geeta Mehta [66]. Hey1, H522, A2780, and MDA-MB-231 were obtained from ATCC. All cell lines were cultured in RPMI-1640 media, supplemented with 10% FBS and 1% Pen Strep, in the cell incubator at 37 °C, with 5% CO2.

### Constructs and transfection

Scramble, MBD3, and CHD4 siRNAs were obtained from Qiagen (1027280, SI00024556, SI00024563, SI00057862, SI04439848). Lipofectamine 3000 transfection reagent (Invitrogen Cat. L3000001) was used to transfect siRNAs in OvCa cells. For each well in a 6-well plate, 5µl of lipofectamine 3000 were used to transfect 60nM indicated siRNA. Media was changed 6 hours after transfection, and cells were collected at 72 hours after transfection for analysis. Scramble, MBD3, and CHD4 shRNAs were purchased from Sigma (T SHC016, RCN0000016153, TRCN0000016154, TRCN0000021361, TRCN0000021363). All shRNAs were stably transduced in OvCa cells by lentivirus.

### Quantitative polymerase chain reaction

RNeasy Mini Kit (Qiagen Cat. 74104) was used to extract RNA, and SuperScript III First-Strand Synthesis System (Invitrogen Cat. 18080051) was used to generate cDNA. qPCR was performed using SsoAdvanced Universal SYBR Green Supermix (BIO-RAD Cat. 1725270). Primers used are listed in Table 1.

### Western blotting and antibodies

For validation of downregulation of MBD3 and CHD4 in serum starvation-induced quiescent cells, cells were treated with normal media and serum-withdrawal media for 72 hours. For testing shRNA efficiency, cells were seeded and cultured for 72 hours. Then, they were treated with trypsin and collected. Cells were lysed with RIPA buffer (Thermo Scientific Cat. 89901) supplemented with Halt Protease and Phosphatase Inhibitor Cocktail (Thermo Scientific Cat. 1861281) and followed by sonication. Protein samples were mixed with Laemmli Sample Buffer (BIO-RAD Cat. 1610747) before heating at 95°C for 7 minutes. Samples were run on 4%-12% NuPAGE Bis-Tris Gel (Invitrogen) in NuPAGE MOPS SDS Running Buffer (Invitrogen Cat. NP0001-02) at 120V for 90 minutes and transferred to a 0.45µm polyvinylidene difluoride membrane in NuPAGE Transfer Buffer (Invitrogen Cat. NP00061) at 30V for 90 minutes. SeeBlue Plus2 Pre-stained Protein Standard (Invitrogen Cat. LC5925) was used as marker.

Membranes were blocked in 5% milk in TBST buffer for 1 hour at room temperature and incubated with 1:1000 anti-MBD3 (Abcam, Ab157464), 1:1000 anti-CHD4 (Proteintech, 14173-1-AP), 1:1000 anti-Histone H3 (Proteintech, 17168-1-AP), 1:1000 anti-LC3B (Proteintech, 18725-1-AP), 1:1000 ATG5 (Cell Signaling, 2630), 1:1000 Ku70 (Cell Signaling, 4588) in 5% milk overnight at 4°C. After washing with TBST three times (10 minutes each), membranes were incubated with 1:10,000 anti-rabbit HRP (Cell Signaling, 7074S) for 1 hour at room temperature. After washing with TBST three times (10 minutes each), Pierce ECL Western Blotting Substrate (Thermo, 32106) and SuperSignal West Femto Maximum Sensitivity Substarte (Thermo, 34095) were used.

### Pharmacological treatments

Vorinostat (S1047), mocetinostat (S1122), belinostat (S1085), carfilzomib (S2853), tanespimycin (S1141), hydroxychloroquine, pidnarulex (S2684), and seclidemstat (S6722) were purchased form Selleckchem. Chloroquine diphosphate salt (C6628-25G) and valproic acid sodium salt (P4543-10G) were purchased from Sigma-Aldrich. Each drug was dose-titrated, and a dose that did not induce cell death was chosen. (Table 2).

**Table 2.** Proteostasis Altering Drugs Evaluated.

| **Name** | **Description** | **Dose** |
| --- | --- | --- |
| **Tanespimycin** | HSP90 inhibitor [70] | 30nM |
| **Carfilzomib** | Proteasome inhibitor [71] | 20nM in PT340 and PT412<br>10nM in Hey1 |
| **Hydroxychloroquine** | Increase pH within intracellular vacuoles, affect the function of lysosome, endosomes, and Golgi apparatus [72] | 50uM in PT340<br>30um in PT412 and Hey1 |
| **Pidnarulex** | rRNA synthesis inhibitor [73] | 150nM |
| **Seclidemstat** | lysine-specific demethylase 1 (LSD1) inhibitor [74] | 300nM |

### Cell proliferation-cell counts

Cell counts were performed by using the Brightfield app on a CellDrop automated cell counter (Denovix).

### Apoptosis assay and flow cytometry

Cells were cultured with indicated conditions for 48 or 72 hours. Floating cells in cultured media and attached cells were collected and mixed. Using BD Pharmingen FITC Annexin V Apoptosis Detection Kit I (BD Biosciences, 556547), cells were stained with Annexin V-FITC and propidium iodide (PI)-PE. CytoFLEX flow cytometer (Beckman-Coulter) and FlowJo were used to analyze samples (20,000 events per sample).

### CellTrace violet staining

PT412 were labeled with CellTrace Violet (Thermo, C34557) as per the manufacturer’s protocol and cultured for 7 days. Flow cytometry was then used to identify CTV-bright and CTV-dim cells.

### Fucci system

Hey1-Fucci cells expressed p27K^-^-mVenus and CDT1(30/120)-mcherry. After DMSO, 3µM vorinostat, 1µM belinostat, 1µM mocetnostat, and 3mM VPA treatment for 48 hours, flow cytometry was performed.

### Multiplexed immunofluorescence and cell cycle mapping

PT340 were seeded in the 8-well chamber (ibidi, 80841) and treated, with or without 3µM vorinostat, for 48 hours. After treatment, cells were fixed with paraformaldehyde for 15 min at room temperature, followed by permeabilization with 0.1% Triton X-100 in PBS for 10 min and blocking in 10% (w/v) donkey serum and 3% (w/v) bovine serum albumin (BSA) in PBS for 1 hour at room temperature.

Primary antibodies were diluted in blocking solution and incubated for 1 hour at room temperature (anti-c-Myc, Abcam, ab190560; anti-p16, Abcam, ab192054; anti-cyclin E2, Abcam, ab207336; anti-phospho- H2AX, Abcam, ab206900; anti-beta-catenin, Millipore, C7738; anti-CDC6, Abcam, ab211734; anti- CDC25C, Abcam, ab205425; anti-CDK2, Bio-Techne, AF4654; anti-CDK4, Abcam, ab213216; anti-CDK6, Abcam, ab198946; anti-cyclin A2, Abcam, ab217731; anti-cyclin B1, Abcam, ab214381; anti-cyclin B2, Abcam, ab250841; anti-cyclin D1, Abcam, ab203448; anti-cyclin D3, Abcam, ab245734; anti-cyclin E1, Abcam, ab194069; anti-E2F1, BioLegend, BL606052; anti-p27, Abcam, ab206927; anti-p53, Abcam, ab224942; anti-phospho-AKT, Cell Signaling Technology (CST), 4075; anti-PCNA, CST, 82968; anti-phospho-ERK, CST, 45899; anti-Plk1, Abcam, ab223901; anti-phospho-RB, CST, 8974; anti-phospho-S6, CST, 3985; anti-phospho-STAT3, CST, 4324). In-house primary antibody fluorescent conjugates were generated using the Alexa Fluor 647 labelling kit (Invitrogen, A20186) or the Alexa Fluor 555 labelling kit (Invitrogen, A20187). Nuclei were labeled with Hoechst (2µg/mL, Invitrogen, H3570) for 10 min at room temperature and imaged in 50% glycerol in PBS. Following imaging, fluorescent dyes were inactivated using an alkaline solution containing H₂O₂ for 15 minutes with agitation, followed by a PBS wash. Samples were re-imaged to measure residual fluorescence.

Subsequent rounds of antibody staining and imaging were then conducted as previously described, starting with primary antibody incubation. Image acquisition, flat-field correction, autofluorescence removal, and registration were performed using the Cell DIVE Imager (Leica Biosystems). Single cells were segmented using Cellpose [24], and intensity measurements were extracted using scikit-image [25]. Feature selection was performed as previously described [26], and cell cycle maps were generated using PHATE [27] as previously described [28].

### RNA sequencing

PT340 and PT412 were treated with or without 3µM vorinostat for 48 hours, and RNA was extracted as described before. The following processes were performed as we described [65]. R studio (2023.12.0+369) and R packages ggplot2 (3.4.4), pheatmap (1.0.12), biomaRT (2.58.2), clusterProfiler (4.10.0), org.Hs.eg.db (3.18.0), and AnnotationDbi (1.64.1) were used for gene oncology (GO) biological process analysis and generating heatmaps and dot plots. To find yeast gene orthologs in humans, g:Orth in g:Profiler was used.[67] Venn diagram was produced by jvenn.[29]

### Breast cancer organoid and CellTiter-Glo 3D cell viability assay

Two triple-negative breast cancer patient-derived organoid (PDO) lines were used. IPM-BO-085, which is Primary TNBC (“TNBC, neoadj AC/T, carbo, atezolizumab vs placebo”) from TP20-M196 (right breast tumor – age 45-50), and IPM-BO-139 (P10, Day 13) – Primary TNBC (“TNBC who progressed through chemo which was aborted”) from TP21-M326 (left breast tumor – age 55-60).

Cells were seeded and cultured 3 days before treatment to allow PDO reformation. Then, organoids were treated with 3µM vorinostat, 20nM carfilzomib, and combination for 3 days. CellTiter-Glo 3D cell viability assays (Promega, G9681) were performed as per manufacture’s protocol.

### Proteasome activity assay

Proteasome activity was measured by monitoring the synthesis of 7-amino-4-methylcoumarin from the Suc-LLVY-AMC as previously described [36] with minor modifications. In brief, PT340-Bright, and PT340-Dim cells were lysed by sonication in ice cold 50 mM Tris-HCl pH 7.5, 1 mM DTT, 0.25 M sucrose, 5 mM MgCl2, 0.5 mM EDTA, and 2 mM ATP. The lysate was mixed with 50 µM N-SUC-LEU-LEU-VAL-TYR-7-Amido-4-methylcoumarin (Sigma S6510) in 20 mM Tris-HCl pH 7.5, 1 mM ATP, 2 mM MgCl2, 0.2% bovine serum albumin, in the presence or absence of 10µM MG-132 (EMD Millipore), a proteasome inhibitor used to confirm the specificity of proteasome activity. The AMC fluorescence (RFU) was measured every 30-60 minutes in a Cytation 5 plate reader (BioTek) at an excitation wavelength of 340 nm and emission wavelength of 420 nm for up to 4 hours.

The proteasome inhibitor MG132 was used to confirm the specificity of proteasome activity.

### Breast cancer liver cell coculture

RFP-labeled MDA-MB-231 cells were seeded onto freshly isolated primary murine hepatocytes as previously described [68, 69]. After 24 hours, cells were treated with DMSO, carfilzomib (10µM), vorinostat (1.5µM), or combination therapy. After 48hours, cells were cultured with 10µM EdU for 24 hours and then fixed with 2% paraformaldehyde at 4°C for 30 minutes. EdU staining was performed using the Click-iT PLUS EdU Alexa Fluor 488 imaging kit (Life Technologies) per manufacturer’s protocol. Total number of RFP+ cells and EdU+ tumor cells was scored with three high-power fields per well [69].

### *In vi*vo model

Female NSG mice, aged 6 weeks, were purchased and given 1 week to acclimate in the animal facility before any procedures were started. The experimental techniques were conducted with the criteria established by the Institute for Laboratory Animal Research of the National Academy of Sciences and were approved by the University of Pittsburgh IACUC.

250,000 cells were injected subcutaneously and bilaterally into the flanks and grew for 5 days before drug treatment. Five days after inoculation, vehicle, 90mg/kg vorinostat, 2mg/kg carfilzomib, combination of 90mg/kg vorinostat and 2mg/kg carfilzomib, 500mg/kg VPA, 30mg/kg chloroquine, and the combination of 500mg/kg VPA and 30mg/kg chloroquine were intraperitoneally injected daily for 3 weeks. Mouse weight and tumor size were measured via calipers and volume calculated using the LxWxW/2 formula.

Tumors were formalin fixed and paraffin embedded. Tumors were cross-sectioned and stained with anti- Ki67. Median sections were scanned and evaluated *in toto* across a minimum of 10 HPF for the three tumor/treatment groups, using ImageJ integrated density and means compared via ANOVA.

### Statistical analysis and software

Statistical analysis was conducted using GraphPad Prism (10.1.1). All data were analyzed using two-tailed Student t tests. A minimum of three replicate experiments (n ≥ 3) were used for statistical analysis. “*” represented p<0.05, “**” represented p<0.01, “***” represented p<0.001, and “****” represented p<0.0001. Data were plotted with means and standard deviations. Synergy calculation was performed by Combenefit using the HSA model [35]. Heatmaps and dot plots were generated by R.

## Acknowledgements

This work was supported by NCI grant 5R01CA278100. RJB is supported by NCI grant P50CA159981. Cancer center core facilities used were supported by P30CA047904.

## Data Availability Statement

RNA-Seq data are in the process of being submitted to the NCI GEO database and should be available at the time of final submission.

## Author Contribution

**Data acquisition:** QJ, ME, AA, SS, JGF, DB**, Data Analysis:** QJ, ME, AA, SS, JGF, DB, JB, RJB, WS, AC, AL. **Bioinformatics:** AS, HUO**. Manuscript Prep.** QJ, AA, ME. **Manuscript editing:** WS, JB, AC, AL, HUO, RJB. **Funding:** AL, JB, AC, RJB.

